# Investigating the long-term stability of protein immunogen(s) for whole recombinant yeast-based vaccines

**DOI:** 10.1101/339093

**Authors:** Ravinder Kumar

**Affiliations:** Section of Molecular Biology, Division of Biological Sciences, University of California San Diego, 9500 Gilman Drive, La Jolla, San Diego, California-92093, USA

**Author notes:** **Correspondence address** Section of Molecular Biology, Division of Biological Sciences, University of California San Diego, 9500 Gilman Drive, La Jolla, San Diego, California-92093, USA.

**Keywords:** Antigen, Stationary-phase, Vaccine, Yeast

## Abstract

Even today vaccine(s) remains a mainstay in combating infectious diseases. Many yeast-based vaccines are currently in different phases of clinical trials. Despite the encouraging results of whole recombinant yeast (WRY) and yeast display (YD), the systematic study assessing the long-term stability of protein antigen(s) in yeast cells is still missing. Therefore, in the present study, I investigate the stability of heterologous protein antigen in the cellular environment of *S. cerevisiae* through *E. coli* surface protein (major curlin or CsgA). Present biochemical data showed that the stationary phase yeast cells were able to keep the antigen stable for almost one year when stored at 2-8 °C and 23-25 °C. Further, iTRAQ based quantitative proteomics of yeast whole cell lysate showed that the level of heterologous fusion protein was low in cells stored at 23-25 °C compared to those at 2-8 °C. In the end, I also proposed a workable strategy to test integrity or completeness of heterologous protein in the yeast cell. I believe that the observations made in the present study will be really encouraging for those interested in the development of a whole recombinant yeast-based vaccine(s).

## Introduction

Vaccines remains an important and mainstay in preventing infectious diseases (Bachler et al. 2013; Perrie et al. 2007; Black et al. 2010). Most of the presently licensed vaccine(s) involve deliberate administration of attenuated or killed pathogen (Shams 2005). Conventional vaccine development regime involved mass culture of an associated or closely related organism and this hampered vaccine development against important infectious diseases including malaria and leprosy (Scollard et al. 2006; Cox 1991). Apart from that, conventional vaccines suffered from various issues neatly summarized elsewhere (Pastoret 1999; Narasimhan et al. 2015; Fang et al. 2016; Lorry et al. 2014).

The arrival of recombinant DNA technology and a parallel improvement in protein purification chemistry allows expression and purification of an unlimited amount of heterologous protein(s) in almost all types of cells ranging from prokaryotes (example *E. coli*, *B. subtilis*) to eukaryotes including yeast, insects, plants, animals. But professional working in concerned discipline chooses yeast as a model for expression of heterologous proteins for use in vaccine development for reasons mentioned elsewhere (Walker 1998; Gellissen et al. 1997; Rose et al. 1987; Strathern et al. 1982; Broach et al. 1991; Valenzuela et al. 1982). So far, yeast (mainly *S. cerevisiae* and *P. pastoris*) emerges as the main workhorse for expression and purification of heterologous proteins with pharmaceutical value. As a result, many of the proteins expressed and purified from yeast are under different phase of clinical trials (Weidang et al. 2014; Ardiani et al. 2010; Bilusic et al. 2014), and protein purified from yeast for the purpose of prophylactic vaccine (example hepatitis B vaccine) is already in market (Valenzuela et al. 1982).

Although peptide-based vaccines somehow overcome the issue associated with the conventional regime of vaccine development and application. But the problem of poor immunogenic response, fast body clearance, an addition of adjuvant and protein stabilizer means a better alternative of both conventional and peptide-based vaccines is must (Purcell et al. 2007; Aguilar et al. 2007). Apart from that, continuous maintenance of 2-8 °C cold chain from point of manufacturing unit till endpoint user especially in resource-poor settings is a big challenge in presently available vaccines (Chen et al. 2011, Das 2004).

Issues associated with conventional and peptide-based vaccine lead to the development of a novel strategy of using yeast in vaccine development which culminates into the use of whole recombinant yeast (WRY) and yeast display (YD). Use of WRY and YD proved much better as these strategies do not involve protein purification, the addition of protein stabilizers and adjuvant (as yeast cell wall acts as natural adjuvant), particulate nature of yeast cells. Apart from that, yeast cells are efficiently taken-up by antigen presenting cells (APCs) including macrophages (Stubbs et al. 2001) and T cells (King et al. 2014, King et al. 2016). Thus, use of WRY and YD appears very promising and, in several cases, WRY based vaccine(s) have already reached various stages of clinical trials. For example, heat-killed whole recombinant budding yeast-based vaccine (GS-4774) against hepatitis B reached to phase-II of clinical trials (Lok et al. 2016) while dead whole yeast was able to protect mice from pulmonary mucormycosis treated with diabetic ketoacidotic-steroid (Luo et al. 2014). Further, injecting killed recombinant yeast expressing hepatitis B virus protein was found safe and well tolerated in healthy subjects (Gaggar et al. 2014) and an oral vaccine against candidiasis was introduced using molecular display systems with *S. cerevisiae* (Shibasaki et al. 2016). All these and similar studies from different labs across the globe clearly showed the merits and benefits of using WRY and YD as yeast-based vaccines over conventional or peptide-based vaccines.

Despite the positive results from different labs against different infectious diseases and cancer a systematic study investigating long-term stability of protein antigens or immunogen in WRY is missing. This compelled us to investigate the long-term stability of heterologous proteins or antigens in the cellular environment of budding yeast. Our present data showed that protein antigens remain stable in yeast cells for up to a year when kept under refrigeration (i.e. at 2-8 °C) and room temperature (i.e. at 23-25°C). We further compared the level of heterologous protein antigens in stationary phased yeast cells kept at 2-8 °C and 23-25 °C and present data showed that yeast cells kept under refrigeration condition were better in keeping the protein antigens. Western blot data was further confirmed by iTRAQ based quantitative proteomics which also showed a differential abundance of CsgA in whole cell lysates from cells stored at room temperature and under refrigerated condition. Surprisingly a minor fraction of cells remains viable even after a year in absence of nutrients, which means that it is important to use yeast strain which can enter into stationary phase, can remain intact but did not grow when came into a nutrient-rich niche like host body.

## Material and methods

### Strains, media and culture condition

Haploid wild-type budding yeast strain of BY4742 background (purchased from EUROSCARF, acc. no. Y10000) expressing *E. coli* CsgA-GFP (fusion protein) under GAP or Glyceraldehydes-3-phosphate dehydrogenase was used in the entire study. Primers used in amplification of curlin major from *E. coli* are (forward primer, PR1) ATGCGAATTCATGAAACTTTTAAAAGTAGCAGCAATTGC, (reverse primer, PR2) ATGCGGTACCGTACTGATGAGCGGTCGCG and (GFP reverse primer, PR3) CGTCGCCGTCCAGCTCGACCAG, used in sequencing. Yeast genetic manipulations were performed as described elsewhere (Longtine et al. 1998; Janke et al. 2004; Güldener et al. 1996). For amplification of CSGA, *E. coli K-12* strain was purchased from ATCC (strain SMG123). GFP was amplified from plasmid purchased from addgene (plasmid # 21052, pESC-URA-ub-G76A-Rnq1-GFP).

Stationary phase was induced and checked in yeast cells as explained previously (Martinez et al. 2004; Kumar et al. 2016) and briefly described here. A single colony of a haploid strain expressing CsgA-GFP (*E. coli* surface protein) under constitutive (GAP or Glyceraldehydes-3-phosphate dehydrogenase) promoter was inoculated in 5 mL YPD (1 % yeast extract, 2 % peptone and 2 % dextrose) and the tube was incubated at 30 °C, 250 rpm for overnight growth. Next day, overnight grown culture was used to inoculate 50 mL YPD such that initial OD_600nm_ was 0.2. Flasks were then incubated at 30 °C; 250 rpm till growth ceased by checking OD_600nm_ of culture by spectrophotometer at regular interval of 24 hour. The culture was regularly checked for any contamination and cell morphology.

### Protein extraction

A known number of cells were re-suspended in TCA (tri chloro acetic acid) such that final concentration of TCA was 20 % and cells were frozen at −80 °C for at least one-hour. After one-hour tubes were taken out of deep the freezer and thawed at room temperature. Tubes were centrifuged at 14000g for 8-10 min and the supernatant was discarded. Resulted pellet was resuspended in 1 mL chilled 100 % acetone using sonicator. Tubes were again centrifuged as above, and the supernatant was again discarded, and protein pellet was air dried and resuspended in protein solubilization buffer (7 M Urea, 2 M Thiourea, 4 % CHAPS) (Reddy et al. 2013).

### Western blots

Western blot was performed as described previously (Kumar et al. 2014).

### Buffer exchange, in solution digestion, iTRAQ labelling, LC-MS/MS, Data acquisition and analysis

For comparing the level of CsgA in yeast cells, quantitative iTRAQ based proteomic analysis was performed. Amount of protein in whole cell lysate was estimated using 2-D Quant Kit (from GE). An equal amount of whole cell lysate protein was labelled with iTRAQ reagent using the 4-plex kit (from AB SCIEX). Rest procedure was performed as described previously. Data acquisition and analysis were performed as explained in detailed elsewhere (Kumar et al. 2016, Reddy et al. 2015). During analysis of MS/MS data using Spectrum Mill (Agilent), taxonomy was kept as *E. coli*.

### Fluorescence microscopy

All the images were captured using Zeiss Axio vision microscope using appropriate filters.

### Accession number

Gene cloned in this study was deposited in Gene bank with following accession number MH264502

## RESULTS

### The basic logic behind the study and workflow

Over the last two to three decades, more than four hundred of papers (including research articles, reviews, case reports) appears which showed the utility of yeast species for expression and purification of heterologous proteins for vaccines (both prophylactic and therapeutic), and other pharmaceutical application. Many of the proteins expressed and purified from yeast are in different phases of clinical trials (Ardiani et al. 2010) and one of them have already hit the markets (Valenzuela et al. 1982) long back.

Further to make the process more rapid, robust and to cut down the cost involved in protein purification, use of whole recombinant yeast and yeast display appeared promising. Many labs already showed the usefulness of whole recombinant yeast and yeast display in development of a yeast-based vaccine(s). But the long-term stability of expressed immunogen (both in whole recombinant yeast and yeast display) remains elusive. Further, a systematic study evaluating the effect of temperature on the long-term stability (here stability means the complete amino acid sequence of the expressed protein) of heterologous proteins is also missing.

Therefore, in the present study, I investigate the stability of heterologous protein *viz* major curlin (*E. coli* surface protein, CsgA-GFP) in the cellular environment of *S. cerevisiae* (with the assumption that cellular environment will be the most suitable environment for the long-term stability of protein). I further study the effect of temperature on the long-term stability of heterologous protein as antigen and the basic workflow of the present study are shown in figure 1.

**Figure 1.**
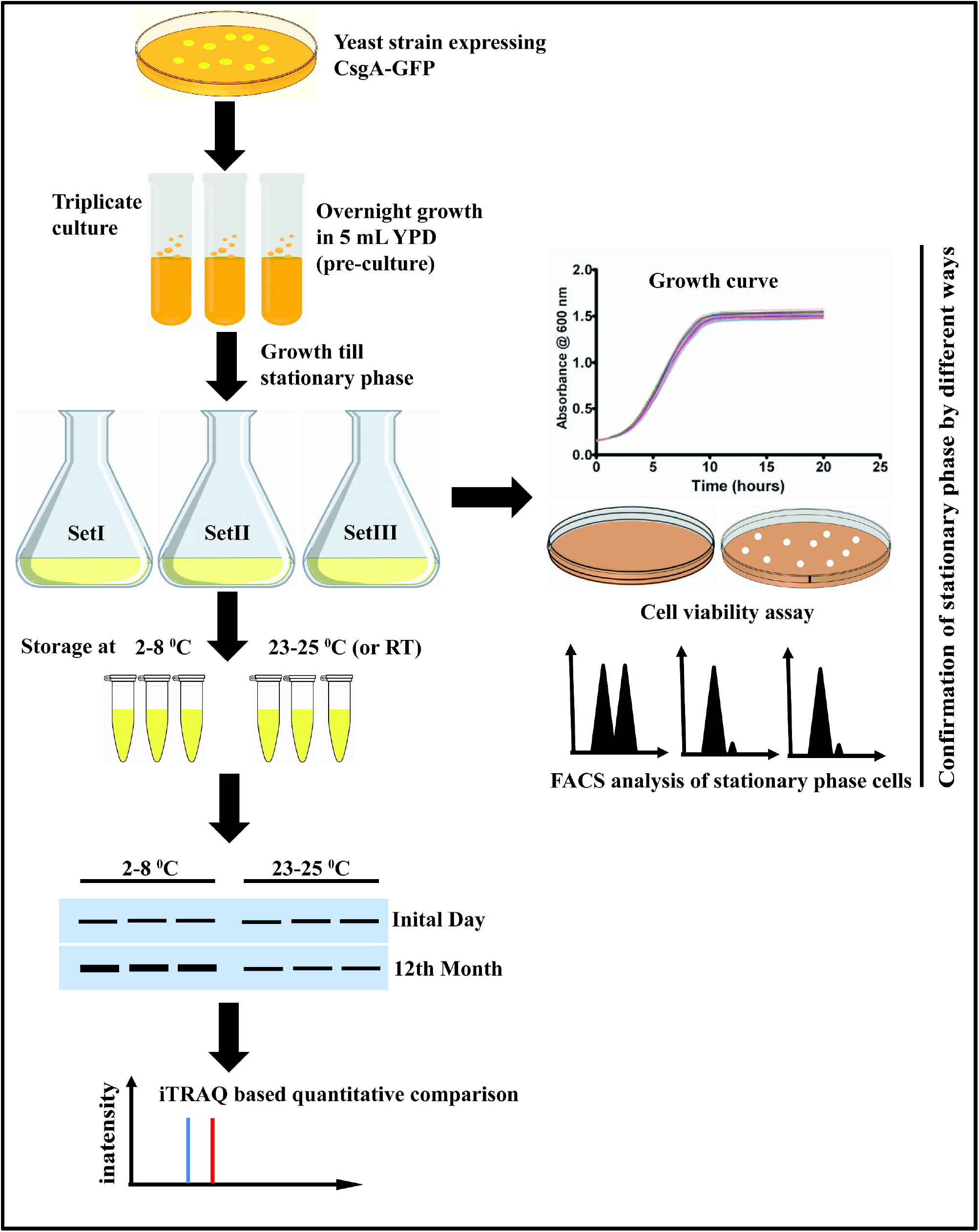
Schematic showing the basic work flow of present study.

The reason for using stationary phase cells are 1) it allows complete utilization of nutrients and harvesting of a large number of cells compared to the culture at mid-log or pre-stationary phase and 2) normal cycling cells cannot be stored at room temperature for a long time let say one year or more and finally 3) stationary phase cells are better in coping the temperature fluctuations which is important in the stability of expressed antigen(s) or immunogen(s).

### Cloning and expression of the CsgA-GFP fusion protein

Although one can express any heterologous protein, just as a proof of concept I am taking bacterial curlin. And believe that other heterologous protein express in yeast will behave similarly from the point of stability in the cellular environment. Curlin is component of bacterial fimbriae, structure important for bacterial attachment to another surface. Homologs of curlin have been already reported in a number of bacterial species and curlin was also detected in surrounding medium (Olsen et al. 1998; Loferer et al. 1997).

*E. coli* surface protein, curlin (CsgA-GFP) was cloned into yeast expression vector under constitutive GAP promoter (for overexpression). Cloning was performed in two steps and maps of both the plasmids (with essential features) used in cloning are shown in fig 2E and fig. 2F for plasmids p1 and p2 respectively, (cartoon presentation) while the nucleotide and amino acid sequence of CsgA are shown in figure 2A and figure 2B respectively. CsgA ORF was PCR amplified from wild-type *E. coli K-12* strain using the primers mentioned in material and methods (Fig 2C). PCR product was treated with *Eco*RI and *Kpn*I restriction endonuclease in CutSmart buffer (from NEB) for one hour and column purified. Parent plasmid with GFP was also treated with same restriction enzymes and gel elution was performed. Ligation was performed at 16 °C for overnight and ligation mix was transformed into *E. coli* competent cells and transformants were selected on LB plus ampicillin plate(s). Positive transformants were checked by sequencing using a reverse primer from GFP. Resulted p1 plasmid (Fig. 2E) was treated with *Eco*RI and *Age*I to release CSGA-GFP cassette (Fig. 2D). Cassette was gel eluted and inserted or cloned into a plasmid having GAP promoter for overexpression. Final plasmid p2 (Fig 2F) was digested with *Stu*I and digested product was transformed into wild-type haploid yeast strain (Güldener et al. 1996). Positive transformants were selected on *his*^−^ plate and expression of a fusion protein (CsgA-GFP) was checked by western blot using anti-GFP antibodies (Fig. 2G). Positive transformants showing proper expression and translated protein (as checked by western blot, Fig 2G) were further used for induction of stationary phase in yeast cells as mentioned in next section.

**Figure 2.**
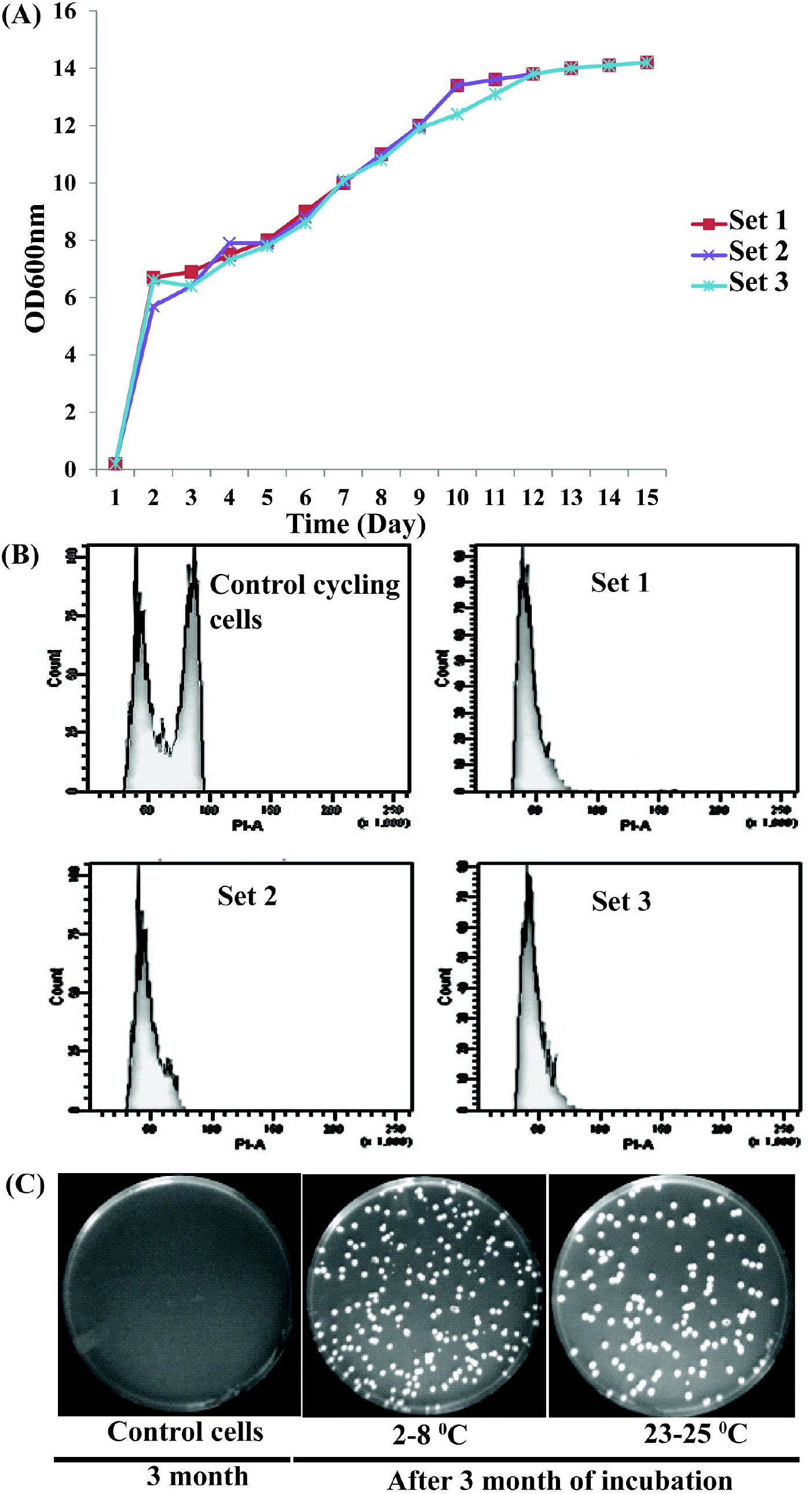
Expression of CsgA-GFP in yeast. (A) The nucleotide sequence of curlin major of *E. coli* along with accession number as given in NCBI. (B) A amino acid sequence of curlin major/CsgA. (C) Agarose gel showing PCR amplification of curlin gene from *E. coli* genome (left panel), (D) release of CsgA-GFP cassette from the p1 plasmid which was introduced in an empty vector having GAP promoter to generate p2 plasmid. Cartoon presentation of map of plasmid (E) p1 and (F) p2 respectively. Note in cartoon presentation only essential elements of plasmids are shown while primers and restriction enzymes used in the construction of these plasmids are mentioned in the text at an appropriate place. (G) Expression of CsgA-GFP from p2 integrated into the genome of wild-type BY4742.

### Induction of stationary phase storage of samples

Positive transformants were patched on fresh YPD plate and plate was incubated at 30 °C till sufficient biomass appears on the plate. These patches were used for inoculating 5 mL YPD and tubes were incubated at 30 °C, 250 rpm for overnight growth. This overnight grown culture was used for inoculation of 50 mL YPD (in 250 mL flask) such that initial OD_600nm_ was close to 0.200. Flasks were incubated at 30 °C, 250 rpm and cell density were checked after every 24 hours (Fig. 3A) till growth ceases and cell entered stationary phase (almost all cells become round) as confirmed by cell morphology (round cells) (Fig. 3B) (Kumar et al. 2016;), cell viability assay (Fig. 3C) as it is known that stationary phase cell can remain viable for extended period even in absence of nutrients (Kumar et al. 2016;) and FACS analysis (Fig. 23) as stationary phase cells arrested at G_1_ (Kumar et al. 2016). Apart from that stationary phase was also checked by detecting Spg4-HA (stationary phase protein 4) which are present in cells only under stationary phase (data not shown) (Kumar et al. 2016). Thus, stationary phase nature of cells was confirmed by different ways before cells were stored for a period of one year.

**Figure 3.**
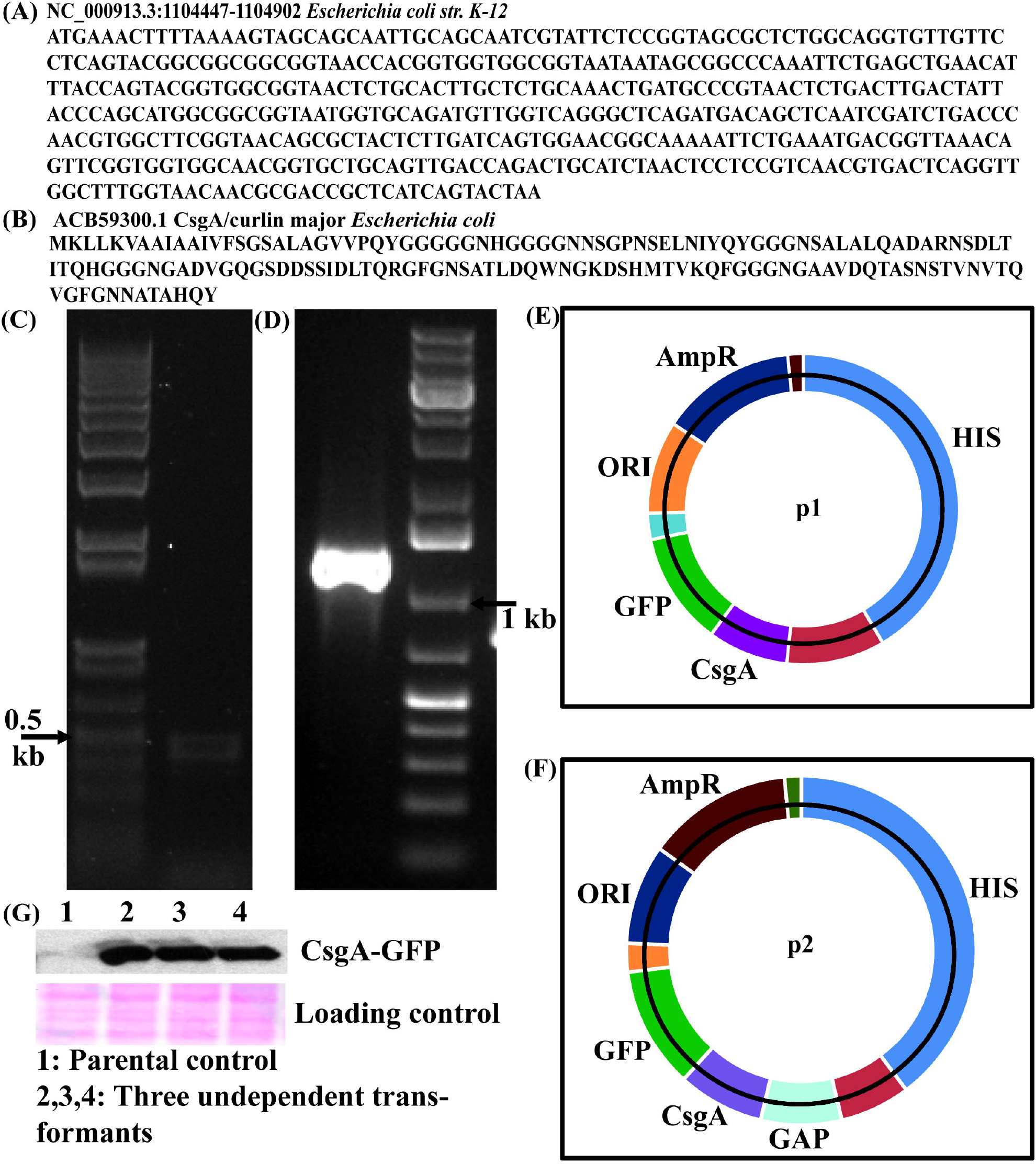
Checking stationary phase in yeast cells. (A) Growth curve of BY4742 expressing CsgA-GFP in three-set (biological triplicate). (B) FACS analysis of cells after 15 days of growth in stationary phase. FACS was performed on all three sets along with control normal cycling cell (overnight culture). (C) Checking the viability of cells after 3 months for both cells stored at 2-8 °C (middle panel), for cells stored at 23-25 °C (right panel) along with control regular cycling cells (overnight grown culture) (left panel).

### Stability of CsgA-GFP fusion protein

From each of three flasks, 1 mL cell suspension was transferred into separate sterile eppendrof tubes such that each tube received around 13 OD_600nm_ of yeast cells. After dispensing cell suspension half tubes were stored at 2-8 °C and remaining half at 23-25 °C. Then after every one month, three tubes each from 2-8 °C and 23-25 °C were taken out and OD_600nm_ were measured till the end of 12 months. Measurement of OD_600nm_ over the year showed that there is a gradual decrease in OD_600nm_ (Fig. 4A), suggesting the cells may be dying. At the end of the one-year final, OD of cells in tubes comes down close to 4.0. In figure, 4A data is shown only for one set, while the experiment was performed thrice, and all three set followed the same trend.

**Figure 4.**
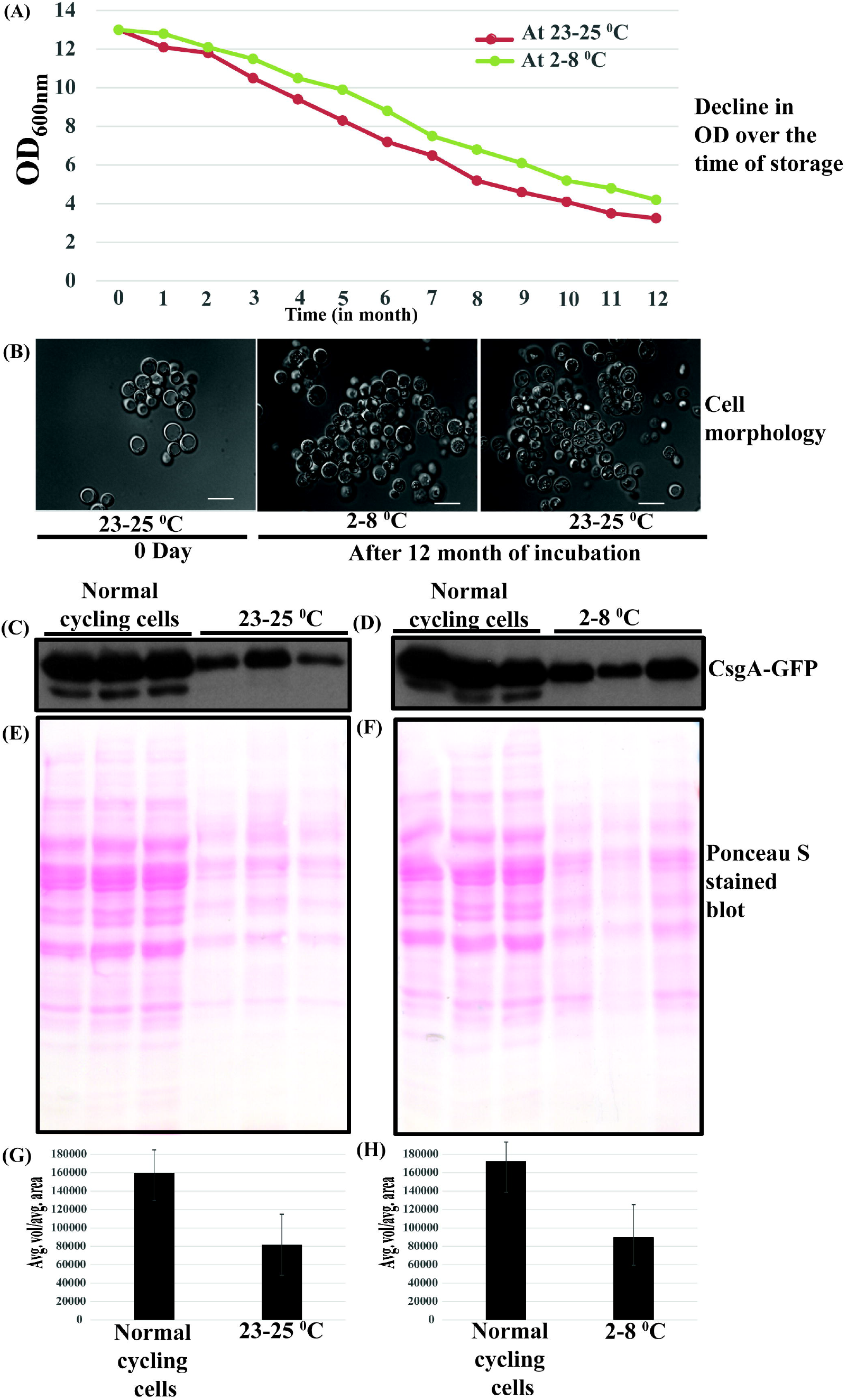
Stability of CsgA-GFP in stationary phase yeast cells. (A) The decrease in OD of cells stored over the period of 12 months under the refrigerated condition and at room temperature. Note experiment was performed in biological triplicate, data is shown for one set, other two set also followed the similar trend. (B) Morphology of cells after one year when stored at 2-8 °C (middle panel), when stored at 23-25 °C (right panel) along with morphology of cells just before storage (left panel). Comparison of the level of CsgA-GFP between equal OD of cells stored at 23-25 °C (C) and stored at 2-8 °C (D) for a duration of one year with that of overnight grown culture. Complete Ponceau S stained blot (E, F) for western blot shown in figure C and D respectively. Histogram showing the intensity of bands (G, H) for blot image C and D respectively.

Further, it was observed that pace with which OD_600nm_ decreases was more in tubes stored at 23-25 °C compared to those stored at 2-8 °C although the difference in OD_600nm_ after one year was not that much significant. Apart from that we also observed the sharp difference in cell morphology after one year (Fig. 4B). The morphology of cells stored under refrigerated condition (Fig. 4B middle panel) was more like to normal cycling cells used as control (Fig. 4B left panel) compared to those stored at room temperature for a year (Fig. 4B right panel). Present microscopic data showed that more number of cells lose their intact nature when stored at room temperature.

After observing and comparing the morphology of cells stored at two different conditions, we check the level of the CsgA-GFP fusion protein in those cells by western blot. Protein in whole the cell lysate was extracted as described in material and methods and proteins were resolved on 10 % SDS-PAGE. Proteins were transferred on to PVDF membrane and the fusion protein was detected using anti-GFP antibodies (Fig. 4C, D). Present western blot data showed that stationary phase yeast cells were able to keep fusion protein (CsgA-GFP or curlin) intact for a period of one year under both the conditions of storage. To give an idea about the nature of whole cell lysate, the complete image of Ponceau S stained blot is also provided below their respective western blot image (Fig. 4E, F for blot image shown in Fig. 4C, D respectively).

The intensity of bands in western blot image in Fig. 4C and D were calculated using Image J software. Histograms showing the combined intensities of the bands in figure 4C, D is shown in figure 4G, H respectively. Thus, combined results of western blot and histogram showed that level of fusion proteins is significantly low when the comparison was made on the number of cells, although heterologous fusion protein remains stable at both the temperature of storage.

I further compare the level of a fusion protein between the cells stored at 2-8 °C to those stored at 23-25 °C and present western blot data suggests that level of a fusion protein is more in cells stored at 2-8 °C compared to those at 223-25 °C (Fig. 5A). Complete blot image for the figure 5A is shown in figure 5B. It is important to mention that western blot data shown so far in figure 4 and figure 5, proteins were normalized based on cell numbers rather on absolute protein quantification. The intensity of bands was calculated using Image J software and histogram showing relative intensities of bands in figure 5A is shown in figure 5C.

**Figure 5.**
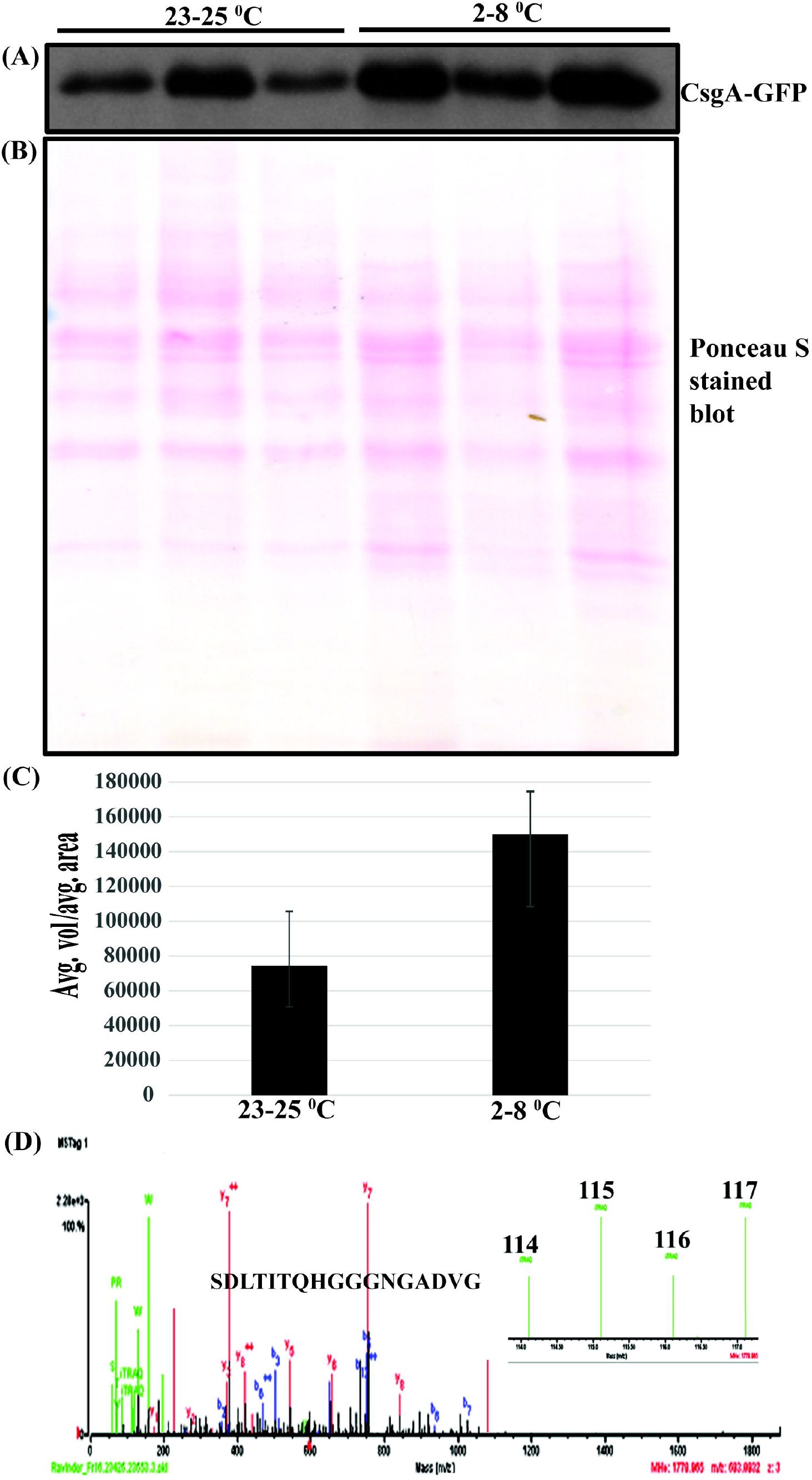
Comparison of CsgA in cells stored at the different temperature. (A) Comparison of the level of CsgA-GFP in cells stored at 23-25 °C (left) and 2-8 °C (right) for a duration of one year. (B) Complete Ponceau stain blot image of western blot shown in figure A. (C) Histograms showing comparative level of CsgA-GFP signal calculated from blot image A. (D) Level of CsgA in cells stored at 23-25 °C (114,116) and those stored at 2-8 °C (label 115,117) as shown by relative intensity of iTRAQ reporter ions. Note in all the western blot protein level was normalized based on the number of cells while in case of iTRAQ, protein in cell lysate was normalized based on protein quantification values.

Further to get a better idea about the relative abundance of a fusion protein in cells stored at two different conditions, the protein concentration in cell lysate was calculated and an equal amount of protein was desalted, digested with trypsin and resulted peptides were labelled with iTRAQ reagents. Present iTRAQ (Ross et al. 2004) based quantitative proteomics showed the level of curlin protein was more in cells stored under refrigerated conditions compared to those which were stored at room temperature (Fig. 5D). Present iTRAQ data also confirmed and validated the western blot data. Just like western blot iTRAQ based quantitative analysis also showed an increased amount of curlin protein (Fig. 5C) in whole cell lysate from cells stored under refrigerated condition compared to those stored at room temperature. In the present study, cell lysate from refrigerated cells was labelled with 115,117 and those at room temperature with 114, 116 labels. It is important to mention that essentially same whole cell lysate was used in both western blots and in the iTRAQ experiment.

I further compare the level of CsgA-GFP using western blot after normalizing protein on basis of protein quantification (like the iTRAQ experiment, mentioned above) and results of western blot are in accordance with iTRAQ data (data not shown).

### Viability of stationary phase cells

Studies from different labs in past and our own previous study (Kumar et al. 2016) showed that stationary phase yeast cells retain the viability for extended periods even in absence of nutrients; therefore I further checked the viability of cells after one year. Cell viability was checked by plating around 300 cells on YPD plates. The platting assay showed that almost all cells lose viability after one year irrespective of the temperature of incubation or storage (Fig. 6A, B). Results of platting assay were further confirmed by checking the viability by trypan blue staining and results of trypan blue staining are in accordance with that of platting assay (Fig. 6C). In another set of experiment, the whole content of tubes (three tubes each from refrigerated condition and from room temperature) was plated on fresh YPD plates and this time we got a huge number of colonies, suggesting a significant number of cells retain viability even after one year in absence of nutrients (data not shown).

**Figure 6.**
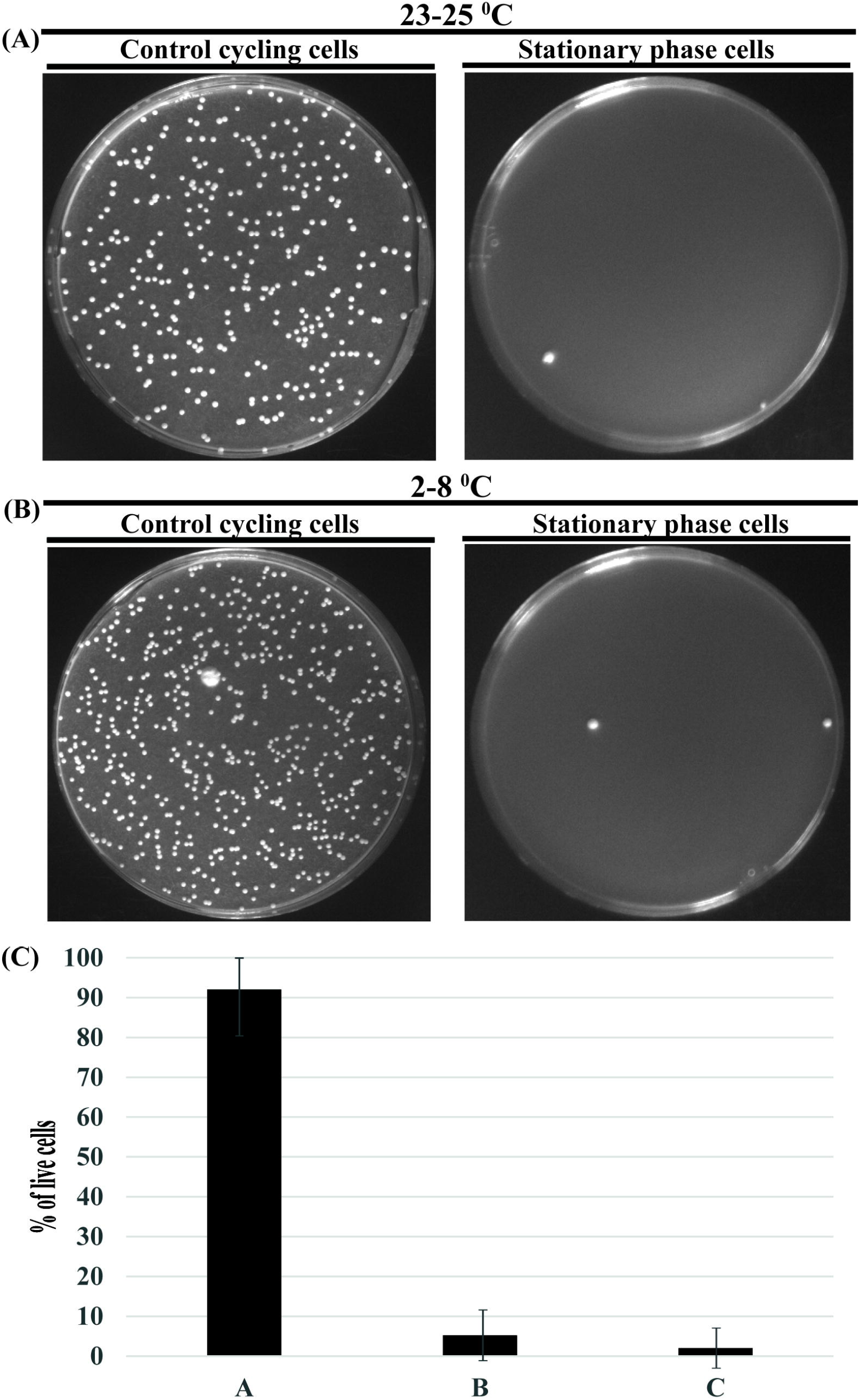
The viability of stationary phase yeast cells. The viability of stationary phased cells after one year of storage (A) at 23-25 °C (right panel) and (B) at 2-8 °C (right panel) compared to control cycling cells (left panel). (C) Cell viability as checked by vital dye trypan blue for both cells stored at 23-25 °C and 2-8 °C compared to control (normal cycling cells).

Although the level of immune response raised by yeast cells expressing immunogen of interest is independent of nature of cells i.e. whether cells are alive or dead (Franzusoff et al. 2005), but still it will not be desirable or advisable to inject live cells especially in individual which are on immunosuppressive medication (example organ transplant) or those with compromised immune system (example AIDs patients). And therefore, it is important that those strains should be used in WRY and YD which can enter into stationary phase, can hold immunogen stable for periods of a year or longer but could not grow when came into the environment rich of nutrients.

## DISCUSSION

In general, peptide-based vaccines overcome many issues associated with the conventional regime of vaccine development as mentioned in the introduction section. But peptide-based vaccines are far away from ideal one and stability of peptide immunogens is an important concern and challenge. Although very few studies are available which assess the effect of temperature on long-term storage of peptide vaccines. For example, a peptide retains its conformation for a week when incubated at 45°C. ESAT651-70-Q11 showed strong immunogenic response even after storage at 45 °C for up to 6 months (Sun et al, 2016). Similarly, in another study, it was observed that mixtures of up to 12 peptides remain stable for up to 5 years when stored at −20 °C or −80 °C (Kimberly et al. 2009). Another study reported that anionic gold nanoparticles and PEG (polyethylene glycol) at the concentration of10^−8^-10^−6^ □M and 10^−7^-10^−4^ □M respectively increased the half-life of a GFP expressing adenovirus from ~48□h to 21 days at 37□° (Maria et al. 2016). All these studies showed that peptide-based vaccine cannot be stored at room temperature for long period and must require deep freeze (−20 °C or −80 °C) which is an important challenge in developing countries. (Chen et al. 2009; Das 2004). Because of this, the introduction of new thermally stable vaccines that do not rely heavily on the cold chain has become an important goal (Chen et al. 2009; Kristensen et al. 2011). Despite this many of the peptide-based vaccines are in different phases of clinical trials (Weidang et al. 2014).

Application of whole recombinant yeast and yeast display clearly overcome many of the limitations associated with conventional and peptide-based vaccines. And many labs have already shown the encouraging results involving WRY or YD (Haller et al. 2005; Schiff et al. 2007; Heery et al. 2015; Hudson et al. 2016; Chaft et al. 2014). All these and similar studies clearly showed the merits of using of whole recombinant yeast-based vaccines. Although several studies are available which investigate the effect of a different temperature on the stability of peptide-based vaccine over a different period of storage. But only one study appeared recently which investigate the amount and the stability of expressed proteins in whole recombinant yeast for a period of six months (Wang et al. 2018). Further, the same study also showed the way for the quantification of heterologous proteins that remain in cells even after a year of incubation can be quantified (Wang et al. 2018).

Still the study which investigates the stability of heterologous protein in WRY for period of year or longer was still missing and therefore this present work tried to fill that important gap in our understanding related to effect of temperature on morphology of yeast cells, stability and level of heterologous proteins in cellular environment of yeast over a period of a year. Although present study showed that the heterologous protein expressed in yeast remain intact but make no comment about the actual amount of antigens that remain in total cell mass even after one year (logic for checking stability is shown through figure 7A). As far as the amount of expressed protein is concerned, incorporating the multi-copy of heterologous genes as described elsewhere (Moon et al. 2016) can take care of this. An important issue which needs to be addressed is whether the level of antigens which remain in yeast cells (over a period of 1-2 years at the different temperature) will be able to raise an optimum immune response. The number of cells that need to be stored at the first hand so that enough amount of antigens remains even after a year of storage for optimum immune response need to be workout. But present study clearly makes a point that WRY based vaccine can be transported and distributed even in absence of cold chain as stationary phase cells are robust enough to keep antigens stable at room temperature for the duration of transportation and distribution which is not the case with the peptide-based vaccine in absence of protein stabilizers.

**Figure 7.**
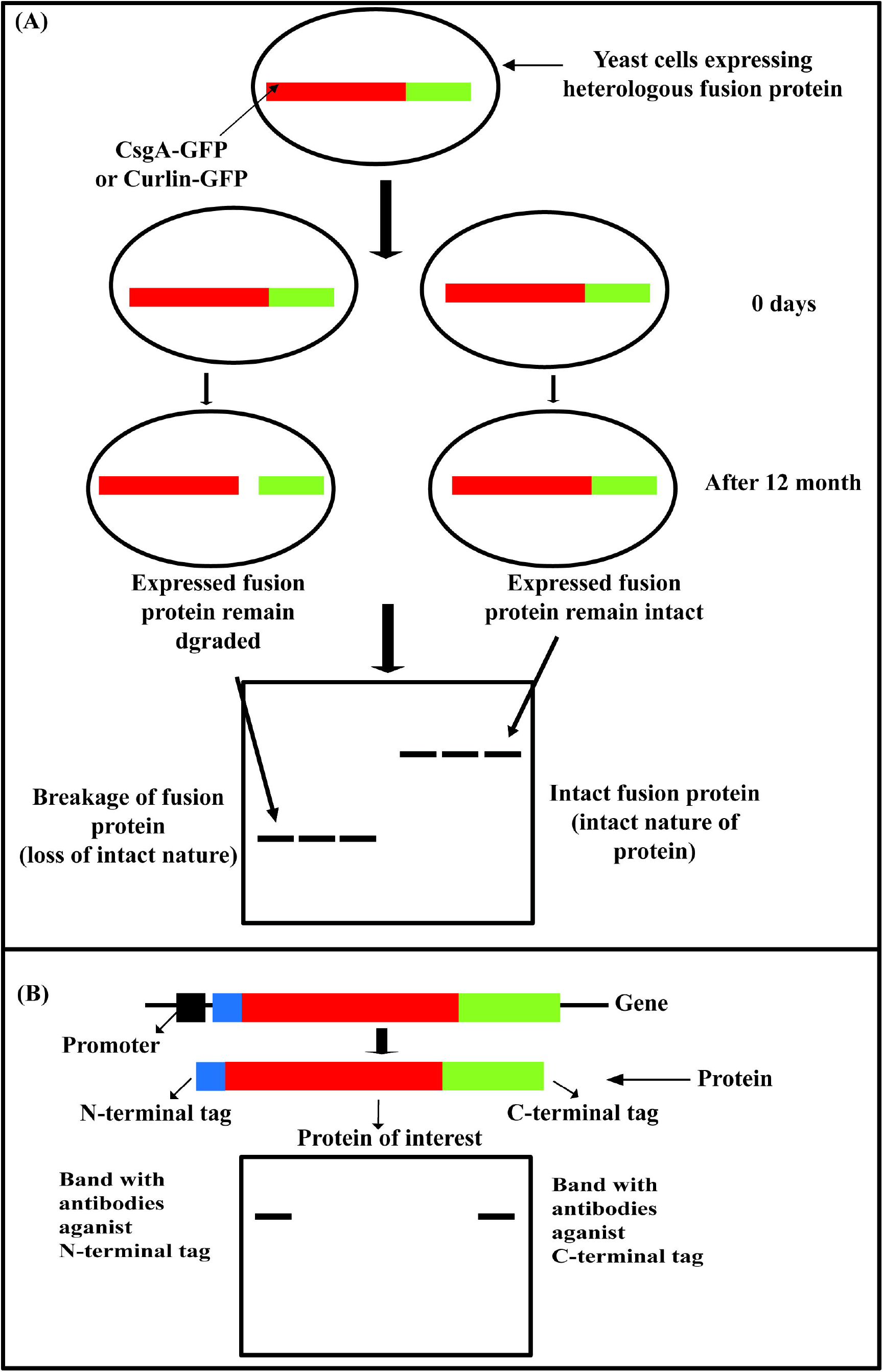
Schematic showing stability of heterologous fusion protein in yeast. (A) Data from present work in which fusion of CsgA and GFP i.e. CsgA-GFP remains intact suggests that expressed heterologous fusion protein remains intact (here intact means complete amino acid sequence without making comment on the conformation of a protein or its structural integrity). (B) Proposed strategy for checking intactness of heterologous protein in whole recombinant yeast using simple western blot by detecting fusion protein using antibodies against N and C terminal tags.

Although the present way of checking the stability of a heterologous protein in WRY is good enough which mainly rely on the intact C-terminal tag and molecular weight markers. But just position of the fusion protein at expected position (relative to protein marker) and one terminal tagging is not the best strategy. Therefore, I also proposed a simple strategy for confirming the completeness of heterologous protein in WRY. This strategy involved tagging of both N and C terminal of heterologous protein with two different tags (Fig. 7B) and detection of the fusion protein with both the tags and at the same place in blot inform that expressed protein is intact from N to C terminus. This strategy was taken from yeast display where the protein of interest is tagged at both N and C terminal (Boder et al. 1997). Bands detected at two different positions when detected using antibodies against N and C terminal tags suggested cleavage or degradation of the heterologous protein.

Since few stationary phase cells remain viable even in absence of nutrients, therefore, it is important to use right yeast strains which are able to enter into stationary phase or G_0_ phase for the long-term stability of immunogen but does not grow on providing nutrients or after administration into subjects. Further, it is known that the magnitude of immune response mounted by the application of whole recombinant yeast is independent of the status of yeast cells i.e. whether cells are viable or dead (Franzusoff et al. 2005). Growing of yeast cells to stationary phase rather than till log or mid-log phase is also advantageous from the commercial point of view as more number of cells or biomass can be generated from given volume of media. This will further help in lower down the cost of the vaccine.

In conclusion, it can be said that soon yeast-based vaccine will be a reality and strategy like WRY and YD will be really crucial in realizing the full potential of the yeast-based vaccine. The cellular environment of yeast appears suitable for keeping the immunogen intact for a long period when stored under a refrigerated condition or at room temperature. Since OD of cells decreases over period storage, it is important to study whether the amount of immunogen in cells will be enough to raise a required immune response. But results of present work will surely encourage more studies in this direction and it will be further interesting to study the stability of expressed immunogen at an elevated temperature close to 37-43 °C which is common in tropical countries of Africa and Asia which require vaccine badly. Again, it will be of utmost importance to finding out the ways to store immunogen in the cellular environment for at least 1-2 years even when cells are stored at elevated temperature (37-43 °C). But present study clearly showed that WRY based vaccine can be transported even under the normal condition without the need for a continuous chain of refrigeration which is itself of great importance.

## Acknowledgement

I am thankful to Dr Piyush Kumar for going through the draft and informing about the required changes in the present manuscript. I am also grateful to UCSD for providing me with the necessary support for completion of this work.

## Authors Contribution

RK conceived, designed, performed the experiments, analyzed the data and wrote the manuscript.

## Compliance with ethical standards

## Ethical approval

This article does not contain any studies with animals performed by any of the authors.

## References

Aguilar JC, Rodriguez EG (2007) Vaccine adjuvants revisited. Vaccine 25:3752–3762.

Ardiani A, Higgins JP, Hodge JW (2010) Vaccines based on whole recombinant *Saccharomyces cerevisiae* cells. FEMS Yeast Res10:1060–9.

Bachler BC, Humbert M, Palikuqi B, Siddappa NB, Lakhashe SK, Robert A Rasmussen, Ruth M Ruprecht. (2013) Novel biopanning strategy to identify epitopes associated with vaccine protection. J. Virol 87:4403–4416.

Bilusic M, Heery CR, Arlen PM, Rauckhorst M, Apelian D, Tsang KY, Tucker JA, Jochems C, Schlom J, Gulley JL, Madan RA (2014) Phase I trial of a recombinant yeast-CEA vaccine (GI-6207) in adults with metastatic CEA-expressing carcinoma. Cancer Immunol Immunother 63:225–234.

Black M, Trent A, Tirrell M, Olive C (2010) Advances in the design and delivery of peptide subunit vaccines with a focus on toll-like receptor agonists. Expert Rev. Vaccines 9:157–173.

Boder ET, Wittrup KD (1997) Yeast surface display for screening combinatorial polypeptide libraries. Nat Biotechnol 15:553–7.

Broach JR, Pringle JR, Jones EW (1991) The molecular and cellular biology of the yeast Saccharomyces. Cold Spring Harbor, NY: Cole Spring Harbor Laboratory Press.

Chaft JE, Litvak A, Arcila ME, Patel P, D’Angelo SP, Krug LM, Rusch V, Mattson A, Coeshott C, Park B, Apelian DM, Kris MG, Azzoli CG (2014) Phase II study of the GI-4000 KRAS vaccine after curative therapy in patients with stage I-III lung adenocarcinoma harboring a KRAS G12C, G12D, or G12V mutation. Clin Lung Cancer 15:405–410.

Chen D, Kristensen D (2009) Opportunities and challenges of developing thermostable vaccines. Expert Rev Vaccines 8:547–557.

Chen X, Fernando GJP, Crichton ML, Flaim C, Yukiko SR, Fairmaid EJ, Corbett HJ, Primiero CA, Ansaldo AB, Frazer IH, Brown LE, Kendall MA (2011) Improving the reach of vaccines to low-resource regions, with a needle-free vaccine delivery device and long-term thermos stabilization. J Controlled Release 152:349–355.

Cox FE (1991) Malaria vaccines-progress and problems. Trends in Biotechnology 9:389–394.

Das P (2004) Revolutionary vaccine technology breaks the cold chain. Lancet Infect Dis 4:719

Fang Y, Liu MQ, Chen L, Zhu ZG, Zhu ZR, Hu Q (2016) Rabies post-exposure prophylaxis for a child with severe allergic reaction to rabies vaccine. Hum Vaccin Immunother 12:1802–1804.

Franzusoff A, Duke RC, King TH, Lu Y, Rodell TC (2005) Yeasts encoding tumour antigens in cancer immunotherapy. Expert Opin Biol Th 5:565–575

Gaggar A, Coeshott C, Apelian D, Rodell T, Armstrong BR, Shen G, Subramanian GM, McHutchison JG (2014) Safety, tolerability and immunogenicity of GS-4774, a hepatitis B virus-specific therapeutic vaccine, in healthy subjects: a randomized study. Vaccine 32:4925–4931.

Gellissen G, Hollenberg CP (1997) Application of yeasts in gene expression studies: a comparison of *Saccharomyces cerevisiae*, *Hansenula polymorpha* and *Kluyveromyce lactis* – a review. Gene 190:87–97.

Güldener U, Heck S, Fielder T, Beinhauer J, Hegemann JH (1996) A new efficient gene disruption cassette for repeated use in budding yeast. Nucleic Acids Res 24:2519–2524.

Haller A, King T, Lu Y, Kemmler C, Gordon G, Bellgrau D, Franzusoff A, Rodell T, Duke R (2005) Whole recombinant yeast-based immunotherapy for treatment of chronic hepatitis C infection induces dose-dependent T cell responses and therapeutic effects without vector neutralization [abstract 132]. Hepatology 42 (Suppl S1):249A.

Heery CR, Singh BH, Rauckhorst M, Marté JL, Donahue RN, Grenga I, Rodell TC, Dahut W, Arlen PM, Madan RA, Schlom J, Gulley JL (2015) Phase I Trial of a Yeast-Based Therapeutic Cancer Vaccine (GI-6301) Targeting the Transcription Factor Brachyury. Cancer Immunol Res 3:1248–1256.

Hudson LE, McDermott CD, Stewart TP, Hudson WH, Rios D, Fasken MB, Corbett AH, Lamb TJ (2016) Characterization of the Probiotic Yeast *Saccharomyces boulardii* in the Healthy Mucosal Immune System. PLoS One 11:e0153351.

Janke C, Magiera MM, Rathfelder N, Taxis C, Reber S, Maekawa H, Moreno-Borchart A, Doenges G, Schwob E, Schiebel E, Knop M (2004) Versatile toolbox for PCR-based tagging of yeast genes: new fluorescent proteins, more markers and promoter substitution cassettes. Yeast 21:947–962.

Kimberly A Chianese-Bullock, Sarah TL, Nicholas ES, John DS, Craig LS Jr (2009) Multi-peptide vaccines vialed as peptide mixtures can be stable reagents for use in peptide-based immune therapies. Vaccine 27:1764–1770.

King TH, Guo Z, Hermreck M, Bellgrau D, Rodell TC (2016) Construction and immunogenicity testing of whole recombinant yeast-based t-cell vaccines. Meth. Mol. Biol 1404:529–545.

King TH, Kemmler CB, Guo Z, Mann D, Lu Y, Coeshott C, Gehring AJ, Bertoletti A, Ho ZZ, Delaney W, Gaggar A, Subramanian GM, McHutchison JS, Shrivastava S, Lee YJ, SKottilil S, Bellgrau D, Rodell T, Apelian D (2014) A whole recombinant yeast-based therapeutic vaccine elicits HBV X, S and Core specific T cells in mice and activates human T cells recognizing epitopes linked to viral clearance. PLoS One 9: e101904.

Kristensen D, Chen D, Cummings R (2011) Vaccine stabilization: Research, commercialization, and potential impact. Vaccine 29:7122–7124.

Kumar R, Dhali S, Srikanth R, Ghosh SK, Srivastava S (2014) Comparative proteomics of mitosis and meiosis in *Saccharomyces cerevisiae*. J Proteomics 109:1–15.

Kumar R, Srivastava S (2016) Quantitative proteomic comparison of stationary/G0 phase cells and tetrads in budding yeast. Sci Rep 6:32031.

Loferer H, Hammar M, Normark S (1997) Availability of the fibre subunit CsgA and the nucleator protein CsgB during assembly of fibronectin-binding curli is limited by the intracellular concentration of the novel lipoprotein CsgG. Mol Microbiol 26:11–23.

Lok AS, Pan CQ, Han SH, Trinh HN, Fessel WJ, Rodell T, Massetto B, Lin L, Gaggar A, Subramanian GM, McHutchison JG, Ferrari C, Lee H, Gordon SC, Gane EJ (2016) Randomized phase II study of GS-4774 as a therapeutic vaccine in virally suppressed patients with chronic hepatitis B. J Hepatol 65:509–516.

Longtine MS, McKenzie A, Demarini DJ, Shah NG, Wach A, Brachat A, Philippsen P, Pringle JR (1998) Additional modules for versatile and economical PCR-based gene deletion and modification in *Saccharomyces cerevisiae*. Yeast 14:953–961.

Lorry G Rubin, Myron J Levin, Per Ljungman, Graham Davies E, Robin Avery, Tomblyn M, Bousvaros A, Dhanireddy S, Sung L, Keyserling H, Kang I (2014) 2013 IDSA Clinical Practice Guideline for Vaccination of the Immunocompromised Host. Clin Infect Dis 58:309–318.

Luo G, Gebremariam T, Clemons KV, Stevens DA, Ibrahim AS (2014) Heat-killed yeast protects diabetic ketoacidotic-steroid treated mice from pulmonary mucormycosis. Vaccine 32:3573–3576.

Maria P, Patrizia A, Jayson P, Marco D’A, Valeria C, Manuela D, Andrea C, Rebecca MB, Nicole H, Paulo JS, Randy PC, Varpu M, Daniel NS, David L, Francesco S, Vincenzo V, Silke K (2016) Additives for vaccine storage to improve thermal stability of adenoviruses from hours to months. Nat. Commun 7:13520.

Martinez MJ, Roy S, Archuletta AB, Wentzell PD, Anna-Arriola SS, Rodriguez AL, Aragon AD, Quiñones GA, Allen C, Werner-Washburne M (2004) Genomic analysis of stationary-phase and exit in *Saccharomyces cerevisiae:* gene expression and identification of novel essential genes. Mol Biol Cell 15:5295–305.

Moon HY, Lee DW, Sim GH, Kim HJ, Hwang JY, Kwon MG, Kang BK, Kim JM, Kang HA (2016) A new set of rDNA-NTS-based multiple integrative cassettes for the development of antibiotic-marker-free recombinant yeasts. J Biotechnol 233:190–199

Narasimhan M, Ahmed PB, Venugopal V, Karthikeyan S, Gnanaraj P, Rajagopalan V (2015) Severe allergic eczematous skin reaction to 2009 (H1N1) influenza vaccine injection. Int J Dermatol 54:1340–1341.

Olsén A, Wick MJ, Mörgelin M, Björck L (1998) Curli, fibrous surface proteins of *Escherichia coli*, interact with major histocompatibility complex class I molecules-Infection and immunity 66:944–949.

Pastoret PP (1999) Veterinary vaccinology. Comptes Rendus De L Academie Des Sciences Serie III-Sciences De La Vie-Life Sciences 322: 967–972.

Perrie Y, Kirby D, Bramwell VW, Mohammed AR (2007) Recent developments in particulate-based vaccines. Recent Pat. Drug Deliv. Formul 1:117–129.

Purcell AW, McCluskey J, Rossjohn J (2007) More than one reason to rethink the use of peptides in vaccine design. Nat. Rev. Drug Discov 6:404–414.

Reddy PJ, Aishwarya AR, Malhotra D, Sharma S, Kumar R, Jain R, Gollapalli K, Pendharkar N, Srikanth R, Srivastava S (2013) A Simple TRIzol Protein Extraction Method For 2-DE, DIGE and MS Analysis of Diverse Samples. Curr. Proteomics 10:298–311.

Reddy PJ, Ray S, Sathe GJ, Gajbhiye A, Prasad TS, Rapole S, Panda D, Srivastava S (2015) A comprehensive proteomic analysis of totarol induced alterations in *Bacillus subtilis* by multipronged quantitative proteomics. J of Proteomics 114:247–262.

Rose AH, Harrison JS (1987) The yeasts. 2nd ed. London/San Diego: Academic Press.

Ross PL, Huang YN, Marchese JN, Williamson B, Parker K, Hattan S, Khainovski N, Pillai S, Dey S, Daniels S, Purkayastha S, Juhasz P, Martin S, Bartlet-Jones M, He F, Jacobson A, Pappin DJ (2004) Multiplexed protein quantitation in *Saccharomyces cerevisiae*using amine-reactive isobaric tagging reagents. Mol. Cell Proteomics 3:1154–1169.

Schiff ER, Everson GT, Tsai N, Bzowej RN, H, G, Gish HG, McHutchison JG (2007) HCV-specific cellular immunity, RNA reductions, and normalization of ALT in chronic HCV subjects after treatment with GI-5005, a yeast-based immunotherapy targeting NS3 and core: a randomized, double-blind, placebo controlled phase 1b study [abstract 1304]. Hepatology 46 (Suppl S1): 816A.

Scollard DM, Adams LB, Gillis TP, Krahenbuhl JL, Truman RW, Williams DL (2006) The Continuing Challenges of Leprosy. Clin Microbiol Rev 9:338–381.

Shams H (2005) Recent developments in veterinary vaccinology. Veterinary Journal 170: 289–299.

Shibasaki S, Ueda M (2016) Oral Vaccine Development by Molecular Display Methods Using Microbial Cells. Methods Mol Biol 1404:497–509.

Strathern JN, Jones EW, Broach JR (1982) The molecular biology of the yeast Saccharomyces: metabolism and gene expression. Cold Spring Harbor, N.Y.: Cold Spring Harbor Laboratory.

Stubbs AC, Martin KS, Coeshott C, Skaates SV, Kuritzkes DR, Bellgrau D, Franzusoff A, Duke RC, Wilson CC (2001) Whole recombinant yeast vaccine activates dendritic cells and elicits protective cell-mediated immunity. Nat. Med 7:625–629.

Sun T, Han H, Hudalla GA, Wen Y, Pompano RR, Collier JH (2016) Thermal stability of self-assembled peptide vaccine materials. Acta Biomater 30:62–71.

Valenzuela P, Medina A, Rutter WJ, Ammerer G, Hall BD (1982) Synthesis and assembly of hepatitis B virus surface antigen particles in yeast. Nature 298:347–350.

Walker GM (1998) Yeast physiology and biotechnology. Chichester, New York: J. Wiley & Sons.

Wang J, Stenzel D, Liu A, Liu D, Brown D, Ambrogelly A (2018) Quantification of a recombinant antigen in an immuno-stimulatory whole yeast cell-based therapeutic vaccine. Anal Biochem 545:65–71.

Weidang Li, Medha D. Joshi, Smita Singhania, Kyle H Ramsey, Ashlesh K. Murthy (2014) Peptide Vaccine: Progress and Challenges. Vaccines (Basel) 2:515–536.

